# Cell Surface Sialoglycan Engineering Through Exogenous mRNA

**DOI:** 10.64898/2026.09.04.749178

**Authors:** Vanessa Affe, Qingyu Shi, Olivia Mann-Delany, David Alvarez, Landon J. Edgar, Haissi Cui

## Abstract

All human cells display a dense matrix of structurally diverse glycans that often terminate in monosaccharides belonging to the sialic acid family of sugars. While these *sialoglycans* are now recognized as critical regulators of human immunity, there is a lack of technologies that provide precise, transient control over their biosynthesis and presentation on a live cell surface.

Here, we addressed this unmet need by developing an mRNA-based platform for delivery of transcripts coding for the enzymes that assemble sialoglycans – sialyltransferases – to human cells. We demonstrated that mRNA coding for the α2-6-specific sialyltransferase ST6Gal1 produced active, Golgi-resident enzyme that potentiated levels of α2-6-sialoglycans on the surface of human cell lines. Increased presentation of these sialoglycans was transient and tracked with the degradation kinetics of the delivered mRNA. Importantly, this approach minimally perturbed other classes of glycans and did not broadly alter physiological transcriptional networks beyond those involved in cellular responses to exogenous RNA. Similar on-target effects were observed for the O-linked α2-6-specific sialyltransferase ST6GalNAc4; however, the effects from mRNA coding for the α2-3-specific sialyltransferases ST3Gal1 and ST3Gal4 were more complex.

Taken together, this mRNA platform provides a framework for cellular sialoglycoengineering endeavours, where precise and reversible control over glycocalyx composition is desired.

## Introduction

Glycans are fundamental regulators of human health and are broadly defined as monosaccharides linked together through regio- and stereo-defined covalent bonds.^1,2^ Many of the glycans presented on human cells terminate in *N*-acetylneuraminic acid (NeuAc), an anionic nonose belonging to the sialic acid family of monosaccharides.^1^ These *sialoglycans* mediate important biological processes, including immune recognition, cell-cell communication, and tissue-specific cell adhesion.^3–6^ Dysregulation of sialoglycan biosynthesis has been implicated in pathologies such as cancer,^7^ coronary artery disease,^8^ and autoimmunity.^9^ Despite their broad biological importance, sialoglycans remain challenging to study because – like all glycans – their structures are not encoded in a template. In other words, it is not possible to *directly* genetically reduce or increase a specific glycan structure.

To circumvent this limitation, the enzymes which catalyze glycan biosynthesis can be targeted instead.^10^ This approach can provide indirect control over cellular glycosylation, which can ultimately illuminate glycan structure-function relationships. For example, genetic knock out and knock in of sialyltransferase genes allows for stable modification of the glycans produced by a cell,^11^ but can also lead to compensatory regulation of related, redundant genes over time. Non-genetic approaches, such as metabolic oligosaccharide engineering (MOE), leverage existing glycan biosynthesis pathways to incorporate exogenously supplemented synthetic monosaccharide analogues.^12,13^ While MOE can be a convenient way to introduce unnatural functionalities into glycans, there are few examples of MOE being used to increase or decrease specific native glycan structures.^14^ Related to this, small molecule inhibitors of glycan biosynthesis^15,16^ have been used extensively to decrease glycan production in cells; however, the current toolbox is small and selectivity is limited to classes of glycans (e.g. all sialosides) rather than specific structures.^13^ Finally, there are several examples of recombinant glycosyltransferases being used to modify cell surface glycans.^17,18^ This approach — selective exoenzymatic labelling (SEEL) — circumvents glycan biosynthetic pathways entirely; however, SEEL is restricted by acceptor glycan presentation on a cell surface and the availability of appropriate nucleotide sugar substrates.^19^

While genetic, MOE, small molecule, and SEEL-based approaches each offer unique advantages, they are limited in their ability to either: (1) provide transient control (genetic),^11^ (2) enforce or increase specific glycan epitopes (MOE, small molecule inhibitors),^14^ or (3) act on the physiological range of available acceptor glycans (SEEL).^18,19^ To address these limitations, we have designed a messenger RNA (mRNA)- based system for cell surface sialoglycan engineering. Our approach potentiates expression of specific sialyltransferases using exogenously delivered messenger RNAs (mRNAs) that code for specific sialyltransferases known to produce defined sialoglycan products. While others have deployed sialyltransferase-coding mRNA for biasing the glycosylation of secreted proteins^20^, use of such mRNA tools for glycocalyx engineering is unexplored. We selected mRNA for this application due to its ability to transiently direct cellular protein synthesis and established potential in clinical settings.^21^ Importantly, advances in *in vitro* transcription (IVT) have enabled the rapid, scalable synthesis of protein coding mRNA from a DNA template.^22^ IVT is also compatible with incorporation of chemically modified nucleosides (N1-methylpseudouridine), which results in mRNA with increased translational capacity and reduced immunogenicity.^23,24^ Further, mRNA delivery circumvents the risk of genomic integration in comparison to DNA therapies and permits *in situ* translation of the target mRNA using the cell’s native translational machinery.^25^ This can aid in the correct physiological subcellular localization and accurate post-translational modification of the target protein.^26^ Critically, mRNA expression is both transient and dose-dependent, providing a level of quantitative, precise control over target mRNA levels that is difficult to achieve through genetic glycoengineering alone. Taken together, these properties position mRNA as an exceptionally versatile and programmable platform for controlling sialyltransferase expression and thereby cell surface glycan composition in mammalian cells.

Here, we show that our mRNA platform provides control over levels of select sialoglycan epitopes. We show that delivery of mRNA encoding the sialyltransferase ST6Gal1 led to specific upregulation of α2-6-sialylation on a human cell surface within 24 hours. Cell surface glycan remodeling was found to occur rapidly upon mRNA delivery and was reversible upon RNA degradation. The extent of α2-6-sialylation was directly proportional to mRNA uptake. We expanded this concept to other sialyltransferases using mRNAs encoding the O-glycan-targeting ST3Gal1 and ST6GalNAc4 as well as the N-glycan targeting ST3Gal4. We probed broadly for changes in cell surface glycosylation for each mRNA and found direct and indirect evidence supporting the installation of increased levels of the targeted sialylation pattern. Notably, mRNA encoding α2-6-producing sialyltransferases produced specific sialoglycan products on the cell surface, whereas mRNA encoding α2-3-producing sialyltransferases produced broader effects on the glycome of transfected cells. Combined, these findings position mRNA-based glycoengineering as a useful tool to transiently modify glycosylation in human cell-based system with control over epitope identity and cell surface glycan levels, but careful construct validation is essential to fully characterize the global impact on cellular glycosylation.

## Results

### Delivery of mRNA encoding ST6GAL1 increased α2-6-sialylation

To understand whether we could modulate cell surface glycosylation using mRNA encoded sialyltransferases, we first tested mRNA coding for *ST6GAL1*. We selected ST6Gal1 due to its well-defined glycan product,^27,28^ which is NeuAc linked α2-6 to galactose (Gal) in the context of *N*-acetyllactosamine (LacNAc) – predominantly on N-linked glycans (**Figure 1A**).^29^ Importantly, these α2-6-linked sialoglycan products can be easily detected using a commercially available lectin (*Sambucus nigra* agglutinin, SNA) as a probe.^28^ We designed plasmids encoding human *ST6GAL1* preceded by a T7 promoter for facile mRNA synthesis and amplified the coding region with primers adding a 45 nucleotide (nt) polyA-tail to the 3’ end. The resulting RNA was treated with DNase I to eradicate contaminating template DNA, and purity as well as integrity were assessed by gel electrophoresis (**Supplementary Figure S1A**). Initial experiments were performed in human embryonic kidney (HEK 293T) cells using a commercially available lipid-based transfection reagent (Lipofectamine MessengerMax), as they are straightforward to transfect and their glycocalyx has been previously mapped and manipulated.^11,30–32^

**Figure 1.**
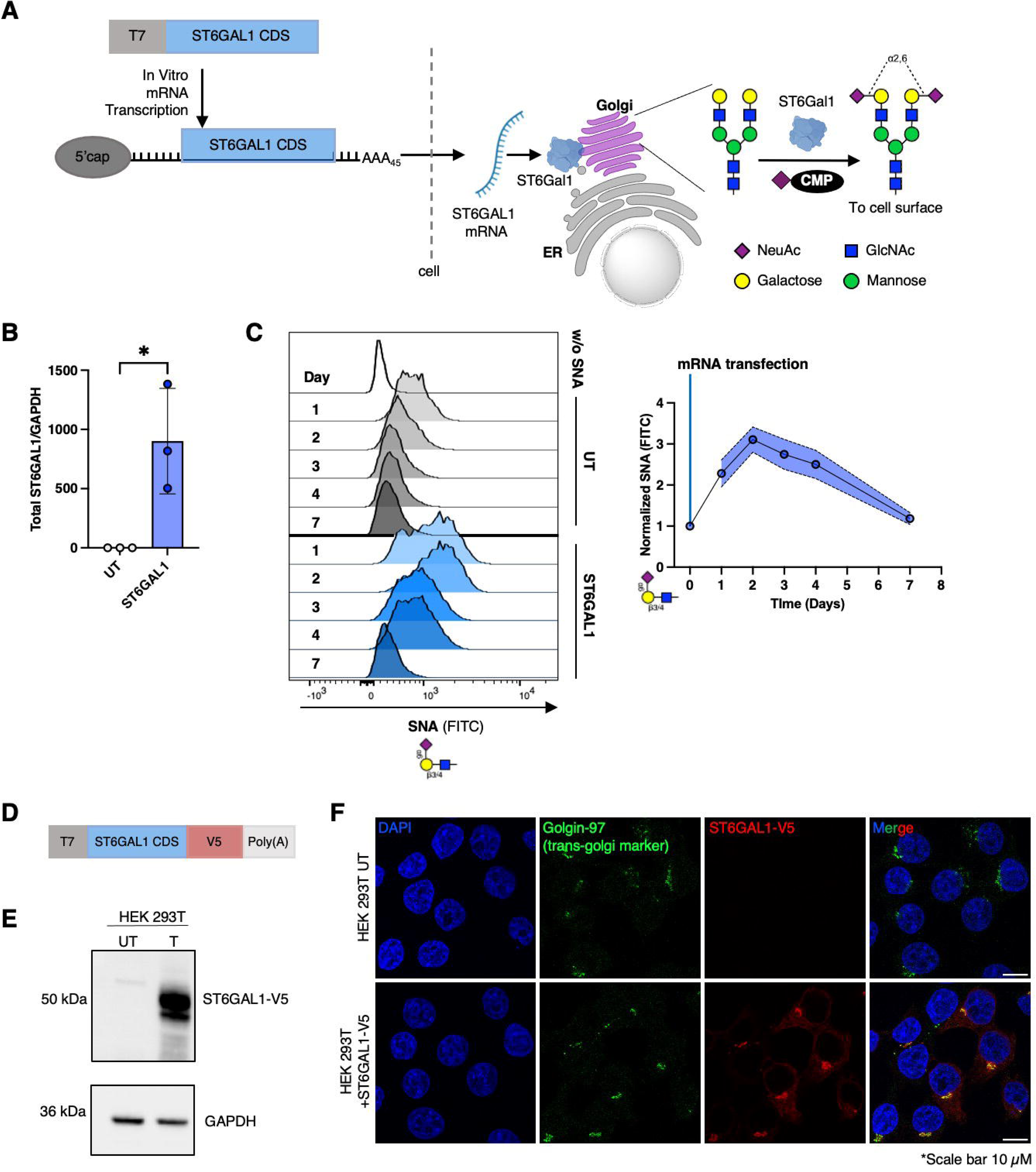
Delivery of *ST6GAL1* mRNA leads to correctly localized protein and elevated 2-6-sialylation on the cell surface. **(A)** Schematic of mRNA coding ST6GAL1 design and expected effect on cell surface sialyation. Galactose (Gal): Yellow circle; Mannose (Man): green circle; N-acetylglucosamine (GlcNAc): Blue square; N-acetylneuraminic acid (NeuAc): Purple diamond. **(B)** qPCR to assess *ST6GAL1* mRNA levels 24 hours after *ST6GAL1* mRNA transfection in HEK 293T cells. Error bars represent mean ± SD (*n =* 3). \**p* ≤ 0.05. Two-tailed unpaired *t*-test. **(C)** *Left panel* — Representative SNA profile of HEK 293T cells in untreated (UT, grey histograms) and *ST6GAL1* mRNA treated (ST6GAL1, blue histogram) cells on days 1, 2, 3, 4, and 7 after transfection, obtained by flow cytometry. *Right panel* — Quantification of SNA intensities from the left panel. SNA data from different time points were normalized to the corresponding untransfected (UT) control. Shaded region represents SD (*n* = 3). **(D)** Schematic of *ST6GAL1-V5* mRNA design. **(E)** Immunoblot confirmation of ST6Gal1-V5 protein expression with anti-V5 antibody 48 hours after *ST6GAL1-V5* mRNA transfection. GAPDH shown as loading control. **(F)** Immunofluorescence-based visualization of Golgin-97 (green) and ST6Gal1-V5 (red) in UT and *ST6GAL1-V5* transfected HEK 293T cells 48 hours after transfection.

We confirmed transfection of HEK 293T cells with *ST6GAL1* mRNA using qPCR (**Figure 1B**). The delivery of exogenous *ST6GAL1* mRNA did not affect endogenous mRNA levels (**Supplementary Figure S1B**). We then monitored α2-6-sialylation using FITC-conjugated SNA over one week via flow cytometry (**Figure 1C**). Importantly, SNA signal increased substantially within 24 h post-transfection, which confirmed that *ST6GAL1* mRNA had a rapid impact on sialoglycan biosynthesis. SNA intensity peaked at 48 h, gradually decreased over the remainder of the experiment, and returned to the level of untransfected control cells after 7 days. This confirmed that *ST6GAL1* mRNA reversibly modified the HEK 293T glycocalyx over a relatively short window of time.

To confirm that the majority of ST6Gal1 protein expressed from exogenous mRNA was trafficked correctly, we generated a variant with a V5-tag to increase the sensitivity of detection and to specifically identify protein products from exogenously introduced mRNA (**Figure 1D**). We confirmed delivery of *ST6GAL1-V5* mRNA via qPCR (**Supplementary Figure S1C**) and as with *ST6GAL1* mRNA, endogenous mRNA levels of *ST6GAL1* remained unchanged (**Supplementary Figure S1D**). The production of ST6Gal1-V5 protein was confirmed through western blotting (**Figure 1E**). To assess ST6Gal1-V5 trafficking to the Golgi apparatus – the compartment where sialyltransferases execute their biosynthetic functions^33^ – we marked the Golgi apparatus with an antibody against Golgin-97, a trans-Golgi associated protein.^34^ We then measured the localization of ST6Gal1-V5 relative to Golgin-97 and found that both proteins were highly co-localized (**Figure 1F**, **Supplementary Figure S1E**), confirming the correct subcellular trafficking of ST6Gal1 produced from exogenously introduced mRNA. In line with this finding, α2-6-sialylation was confirmed to be increased evenly along the surface of *ST6GAL1-V5* transfected HEK 293T cells by immunofluorescence staining with SNA lectin (**Supplementary Figure S1F**).

Our next goal was to assess the correlation between α2-6-sialylation and *ST6GAL1* mRNA uptake. To track mRNA delivery on a single cell basis while preserving compatibility with established lectin staining protocols, we generated a new mRNA template, which encoded both *ST6GAL1* and the fluorescent reporter protein dTomato (ST6GAL1/dT). To avoid perturbation of ST6Gal1 activity through a fusion protein, we opted to integrate a P2A sequence,^35^ which leads to ribosome skipping to the dTomato open reading frame to yield two separate proteins from the same mRNA (**Figure 2A**). This allowed us to bin cells based on dTomato expression – dT^high^ and dT^low^ – which we used as an indirect metric for higher or lower uptake and translation of *ST6GAL1/dT* mRNA, respectively (**Figure 2B**). We again confirmed *ST6GAL1* mRNA delivery using qPCR (**Supplementary Figure S2A**). Of note, the delivery of mRNA encoding dTomato alone did not increase SNA signal, confirming that the effect on the glycocalyx was not simply the result of a general response to mRNA (**Supplementary Figure S2B**).

**Figure 2.**
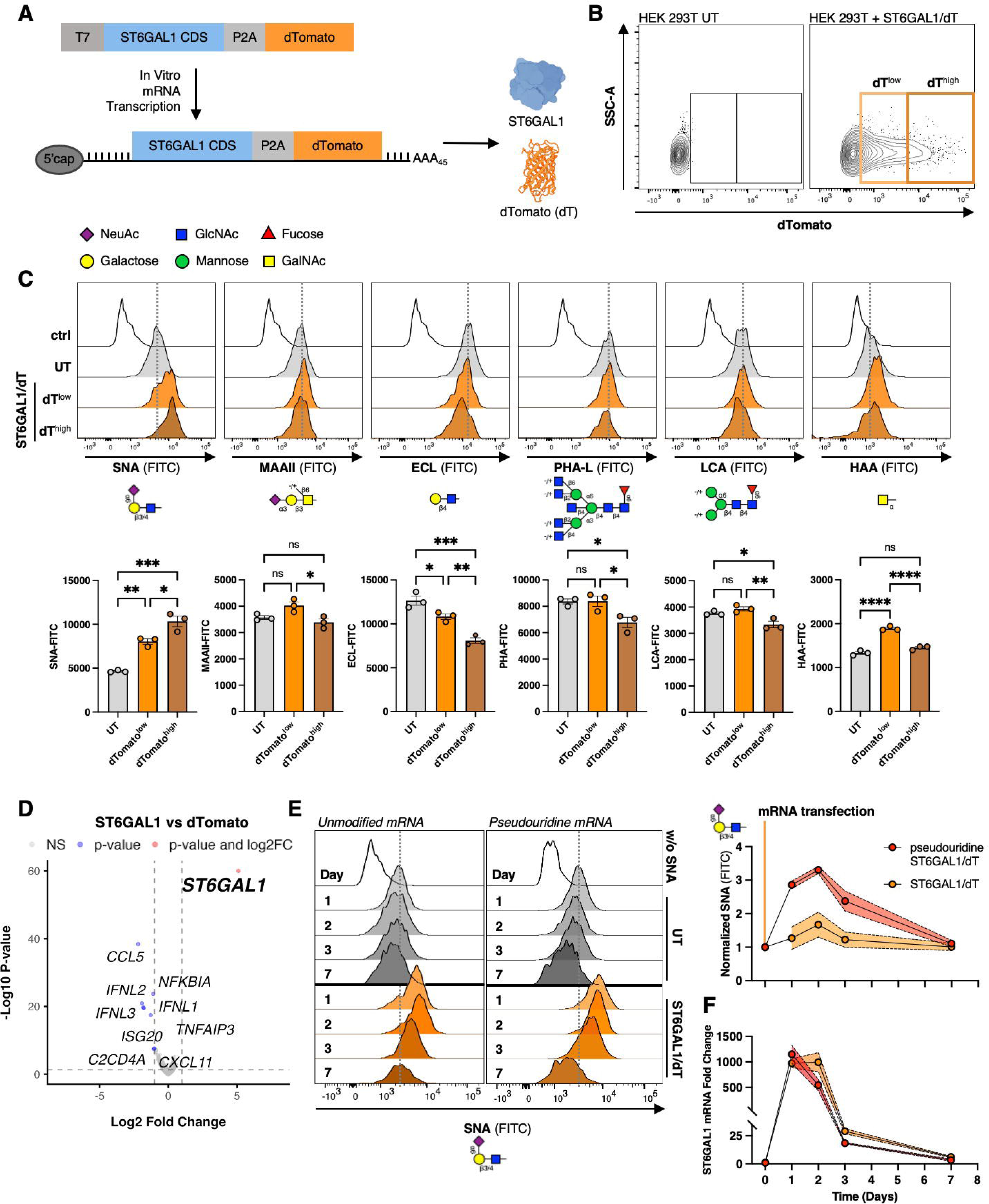
Glycocalyx engineering through delivery of ST6GAL1/dTomato mRNA (*ST6GAL1/dT*) is precise. **(A)** Schematic of *ST6GAL1/dT* mRNA design. **(B)** Gating scheme for flow cytometry analysis based on dTomato expression in HEK 293T cells. Untransfected (UT) control and HEK 293T + *ST6GAL1/dT* cells. dT^low^: cells with lower dTomato expression; dT^high^: cells with higher dTomato expression 48 hours after transfection. **(C)** *Top panel* — Representative glycan profiles of HEK 293T cells 48 hours after *ST6GAL1/dT* mRNA transfection measured by flow cytometry. *Bottom panel* — Quantification of data from Top panel. Error bars represent Mean ± SD (*n* = 3). ns = *p* ≥ 0.05, \**p* ≤ 0.05, \*\**p* ≤ 0.01, \*\*\**p* ≤ 0.001, and \*\*\*\**p* ≤ 0.0001. One-way ANOVA followed by Tukey’s multiple comparisons test. **(D)** RNAseq identified differentially expressed genes in *ST6GAL1* mRNA vs *dTomato* mRNA transfected HEK 293T cells after 48 hours. The X-axis shows the Log_2_ Fold Change (effect size), and the Y-axis shows the –Log₁₀ adjusted p-values (significance). **(E)** *Left panel* — Representative SNA profiles of HEK 293T cells treated with unmodified *ST6GAL1/dT* mRNA (left) and pseudouridine modified *ST6GAL1/dT* mRNA (right) on day 1, 2, 3, and 7 after transfection, measured by flow cytometry. *Right panel* — Quantification of SNA data from left panel. SNA data from different time points are normalized to their corresponding UT sample. Shaded region represents SD (*n* = 3). **(F)** qPCR to assess the degradation kinetics of both unmodified and pseudouridine mRNA normalized to GAPDH on days 1, 2, 3, and 7 after transfection. Shaded region represents SD (*n* = 3).

### Increasing ST6Gal1 has a predictable, specific impact on the glycocalyx composition of ST6GAL1 mRNA-transfected cells

We next assessed changes in the glycocalyx by deploying a panel of six lectins that each detect a unique glycosylation pattern.^27^ Our goal was to assess if changing the production of α2-6-sialoglycans through potentiated ST6Gal1 expression could impact the biosynthesis of other classes of glycans. Consistent with expectations, dT^high^ cells showed the highest amount of SNA staining relative to dT^low^ and control cells and followed a pattern expected for a dose-response relationship (**Figure 2C, Supplementary Figure S2C**). In line with this, staining with *Erythrina Cristagalli* Lectin (ECL), which binds to uncapped (asialo) LacNAc epitopes, was diminished in cells transfected with *ST6GAL1/dT* mRNA. Combined, these data confirmed that as more α2- 6-sialoglycans were produced by ST6Gal1, fewer acceptor LacNAc sites remained. Further, at higher ST6Gal1 levels, binding of the lectin from *Phaseolus vulgaris* (PHA-L), which detects branched N-linked glycans, was decreased. This was in line with known inhibition of PHA-L binding when LacNAcs on branched N-linked glycans terminate in sialic acid,^27^ consistent with the function of ST6Gal1. In contrast, staining with lectin from *Lens culinaris agglutinin* (LCA), which detects (fucosylated) chitobiose-Man_3_, was only subtly affected. We also observed minor changes in *Maackia amurensis* Lectin II (MAAII) and *Helix aspersa* agglutinin (HAA) binding, which detect O-linked α2-3-sialyl core 1 (sialyl T antigen) and O-linked α-*N*-acetylgalactosamine (GalNAc), respectively. These changes were not consistent across cells expressing higher or lower amounts of exogenous ST6Gal1, as indicated by dT fluorescence. Combined, these data suggested that most changes in glycan epitope presentation on *ST6GAL1/dT* mRNA transfected cells reflected the specific installation of additional α2-6-sialoglycans at the expense of uncapped galactose. The overall glycocalyx composition was not severely disrupted beyond these expected changes, confirming the utility of this approach for on-target and predictable glycocalyx engineering.

While these results confirmed the predicted effects of *ST6GAL1/dT* mRNA on α2-6- sialylation, they did not report on global changes to sialoglycan density on the cell surface. To measure this, we leveraged the synthetic monosaccharide per-*O*-acetyl-*N*-azidoacetylmannosamine (Ac_4_ManNAz), which is a click chemistry-compatible probe that is metabolically converted into CMP-*N*-azidoacetyl sialic acid (CMP-SiaNAz) in cells.^12,36^ CMP-SiaNAz can be used as a substrate by all sialyltransferases and therefore reports on global changes to the sialoglycan content of a cell, irrespective of chemical linkage or acceptor glycan identity. We found that treatment of cells with Ac_4_ManNAz before and after *ST6GAL1* mRNA delivery did not lead to detectable changes in sialoglycan density on the cell surface (**Supplementary Figure S2D-E**). This suggested that the overall sialoglycan content of the HEK 293T cells was not substantially perturbed. Instead, we propose that the newly installed sialoglycans were likely produced at the expense of other classes of glycans, such as its acceptor substrate, LacNAc.

We further probed this by measuring if transient *ST6GAL1* overexpression led to adaptive changes in the expression of other glycosyltransferases. We used RNAseq of poly-A enriched RNAs to broadly measure changes in gene expression at the mRNA level. Normalized read counts confirmed a >100-fold overall increase in *ST6GAL1* mRNA (**Supplementary Figure S2F**) 48 hours post-transfection with *ST6GAL1* mRNA, compared to *dTomato*-mRNA transfected control cells. Overall changes in differential expression between *dTomato* control mRNA and *ST6GAL1* mRNA were minor (**Figure 2D, Supplementary Table S1**) and mostly limited to immune response and viral response pathways (**Supplementary Figure S2G, Supplementary Table S2**), as expected for the transfection with unmodified mRNA. Of note, these pathways were downregulated in comparison to cells transfected with a control mRNA. No significant compensatory regulation of other glycosyltransferase genes was observed. This confirmed that *ST6GAL1* mRNA delivery was a strikingly selective approach for the installation of additional α2-6-sialoglycans on the cell surface without interfering with other glycosyltransferases and metabolic pathways.

### Potentiation of mRNA half-life through pseudouridine incorporation increased the magnitude and duration of sialoglycan engineering

The studies described above were conducted using unmodified RNA to take advantage of the short half-life of unmodified mRNA in cells. We next tested whether an extension of mRNA half-life and an increase in mRNA translation would lead to a more durable and pronounced effect on glycosylation. To realize this, we modified the *ST6GAL1/dT* mRNA product with pseudouridine (ψ) (**Figure 2E**, **Supplementary Figure S2H** for gating scheme). Replacement of uridine with ψ increases mRNA translation.^37^ Incorporation of ψ also dampens innate immune responses to exogenous RNA^38^. We found that ψ-modified *ST6GAL1/dT* mRNA produced a more robust increase in α2-6 sialylation vs unmodified RNA, as reported by increased SNA staining: 3.5-fold vs. 1.5-fold relative to control cells, respectively (**Figure 2E**). In addition, the response was prolonged; while cells treated with unmodified RNA returned to baseline levels on day 3, pseudouridine RNA treated cells still displayed an over 2.5-fold increase in SNA levels vs. untransfected controls. The kinetics of α2-6-sialylation were closely correlated with the rate of mRNA degradation in cells transfected with unmodified mRNA (**Figure 2F**). Interestingly, while ψ mRNA prolonged the effect on sialylation, the kinetics of mRNA clearance were similar to unmodified RNA as measured by qPCR. Combined, these data confirmed that ψ-incorporation could increase the intensity and duration of mRNA-mediated sialoglycoengineering. With this established, we opted to continue using unmodified RNA for the remainder of the study to capture transient cell surface glycoengineering, which is a major advantage of our approach.

### mRNA-driven sialoglycan engineering in a human T cell line

Regulation of cell surface glycosylation has been extensively shown to modulate immune cell activation, including on T lymphocytes, which we and others have previously explored.^3,39–42^ To investigate the potential for mRNA-based glycoengineering to control T cell glycosylation, we next performed preliminary studies using an immortalized human T lymphoblast cell line (Jurkat) as a model T cell system. Since these cells already presented substantial α2-6-sialylation (**Supplementary Figure S3A**), we opted to include our previously reported^41^ *ST6GAL1* knock out (*ST6GAL1*^−/−^) Jurkat line as a parallel sample group. We envisioned that *ST6GAL1*^−/−^ cells would be a useful canvas for tracking the kinetics of α2-6-sialylation without interference from endogenously produced glycans. As expected, *ST6GAL1*^−/−^ cells were devoid of SNA staining (**Supplementary Figure S3A**), confirming that ST6Gal1 was the main contributor to Jurkat cell surface α2-6-sialylation.^41^

Our first experiments involved transfection of WT Jurkat cells with *ST6GAL1* mRNA; however, we did not observe significant changes in SNA staining (**Supplementary Figure S3A**). In contrast, *ST6GAL1*^−/−^ counterparts displayed an increase, although the distribution was bimodal. To control for either insufficient transfection efficiency or varying degrees of mRNA uptake between individual cells, we pivoted to the *ST6GAL1/dT* mRNA system (**Supplementary Figure S3B, C**). This allowed us to selectively gate on successfully transfected cells with similar levels of mRNA uptake as reported by dTomato signal (**Figure 3A)**. Here, we found that while a low percentage of WT Jurkat cells expressed dTomato (15%), the number was substantially increased for *ST6GAL1*^−/−^ cells, which showed similar percentages of dTomato positive cells as their HEK 293T counterparts (70%) (**Figures 2B & 3A**). This suggested that a reduction in cell surface sialylation increased transfection efficiency or mRNA translation. Next, we tracked α2-6-sialylation in Jurkat cells over time by again gating on dT^high^ populations and measuring SNA staining (**Figure 3B**). Here, we measured a 1.5–2-fold increase in SNA staining for dTomato positive WT Jurkat cells, which was in line with findings for HEK 293T cells. In *ST6GAL1*^−/−^ cells, exogenous delivery of *ST6GAL1/dT* mRNA restored α2-6-sialylation to near-WT levels, leading to a >6-fold increase in SNA signal vs. untransfected controls (**Figure 3B**). Restoration of α2-6-sialylation on *ST6GAL1*^−/−^ cells was also transient, as it decreased after 3–4 days – this was in line with the cytosolic half-life of unmodified mRNA in HEK 293T cells (**Figure 2F**). Sialoglycan dynamics were faster in Jurkat cells, with SNA levels peaking after 24 hours as compared to 48 hours in HEK 293T cells (**Figures 1C**, **2E**, and **3B**). Taken together, these results suggested that the magnitude of sialoglycan engineering through mRNA differed between cell types but was overall comparable between cell systems.

**Figure 3.**
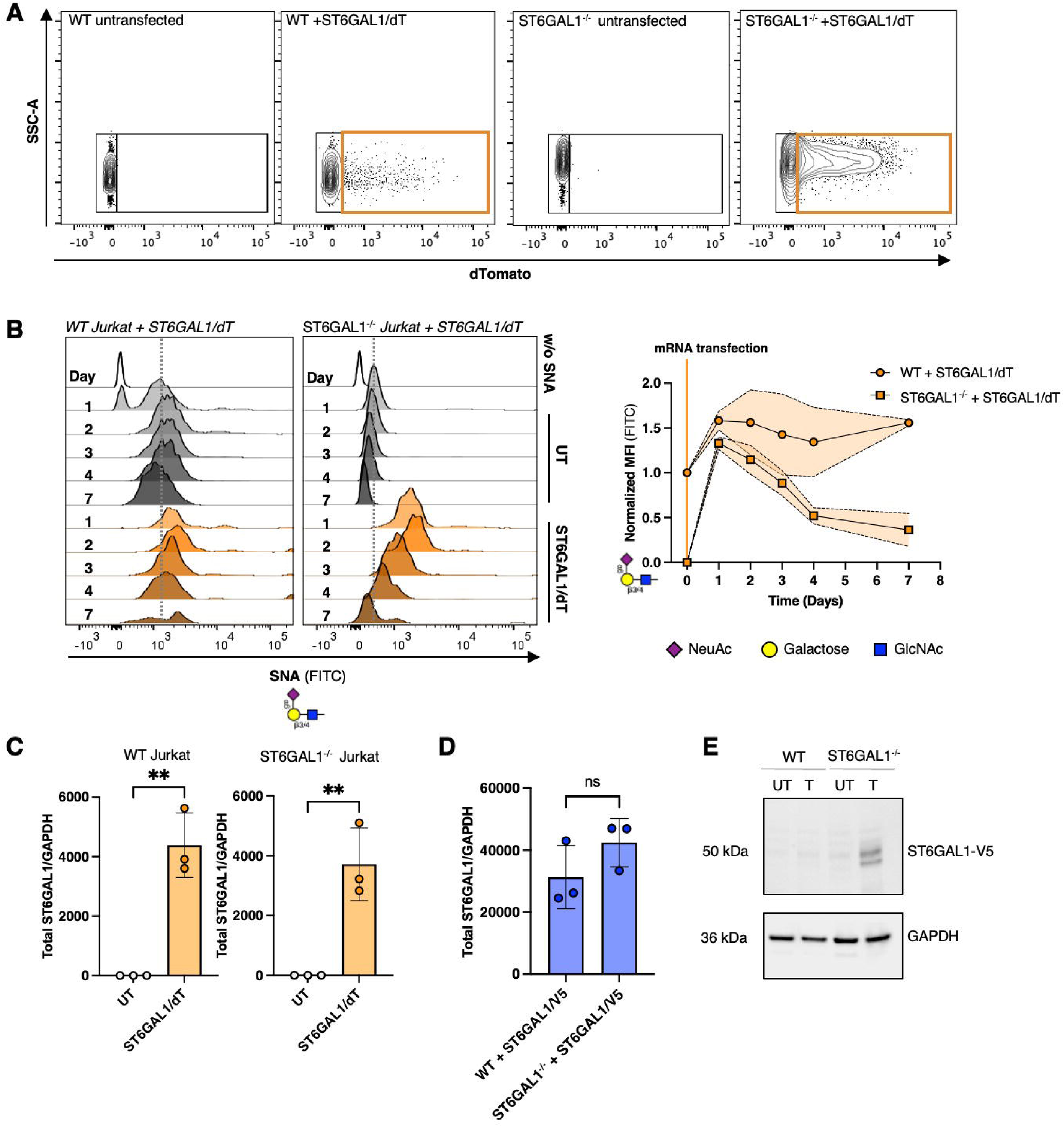
*ST6GAL1/dT* mRNA delivery in a lymphocyte-derived cell line. **(A)** Flow cytometry gating scheme for dTomato+ cells in WT and *ST6GAL1-/-* Jurkat cells. Orange box denotes dTomato positive cells. **(B)** Left panel — Representative histograms of WT and *ST6GAL1-/-* Jurkat cells treated with *ST6GAL1/dT* mRNA on days 1, 2, 3, and 7 after transfection, measured by flow cytometry. Right panel — Quantification of SNA signal from left panel. SNA data from different time points are normalized to untransfected *ST6GAL1-/-* Jurkat cell SNA signal. Shaded region represents SD (n = 3). **(C)** qPCR confirming the delivery of *ST6GAL1/dT* mRNA to WT and *ST6GAL1-/-* Jurkat cells 24 hours after transfection. **(D)** qPCR confirming the delivery of *ST6GAL-V5* mRNA to WT and *ST6GAL1-/-* Jurkat cells 24 hours after transfection. **(E)** Immunoblot to compare protein levels of ST6Gal1-V5 in WT and *ST6GAL1-/-* Jurkat cells 24 hours after transfection, GAPDH shown as loading control. (**C-D)** Dots show biological replicates. Mean ± SD (*n* = 3). ns = *p* ≥ 0.05, \*\**p* ≤ 0.01, Two-tailed unpaired *t*-test.

A comparison of the kinetics of α2-6-sialylation across the WT and *ST6GAL1*^−/−^ Jurkat cells suggested that increases in sialylation were dependent on the base level of glycosylation (**Figure 3B**): even when comparing dTomato^high^ cells, *ST6GAL1*^−/−^ Jurkat cells displayed higher increases in SNA binding after transfection. This difference could also be explained by an intrinsic upper limit to cell surface sialylation in this system. To test if differences in transfection efficiency could explain these results, we compared *ST6GAL1/dT and ST6GAL1-V5* mRNA levels between the Jurkat cell lines following transfection. Here, we found that levels of both exogenously delivered mRNA were comparable in bulk cell lysates (**Figure 3C, D**), despite the discrepancy in the percentage of successfully transfected cells (**Figure 3A**). Next, we assessed ST6Gal1 protein production following transfection with *ST6GAL1-V5* mRNA and found that while *ST6GAL1*^−/−^ cells produced detectable amounts of the protein product (ST6Gal1-V5), WT counterparts did not (**Figure 3E**). An explanation for this could be that the expression of the ST6Gal1 protein itself was capped and that protein synthesis or stability were regulatory handles for controlling sialoglycan levels on a cell surface. Alternatively, transfection might not have reached sufficient cells for detectable protein production in bulk. These results support the need for optimization of mRNA delivery to cell types that are challenging to transfect and lend motivation for further studies into whether sialyltransferase production is intrinsically capped.

### Effects of ST6GAL1-driven glycocalyx remodelling across different model systems

We next compared broader changes in cell surface glycosylation across our model systems to further evaluate the selectivity of *ST6GAL1* mRNA for glycoengineering applications (**Figure 4A**, **Supplementary Figure S3D, E**). As expected, we detected only minor changes in MAA-II and LCA binding in *ST6GAL1/dT*-positive cells across all three model cell systems. Decreases in ECL and PHA-L staining were also observed across all cell lines, consistent with capping of acceptor LacNAc sites and loss of PHA-L affinity following sialylation of branched N-linked glycans. Surprisingly, we observed a small, albeit significant, increase in O-linked *N*-acetylgalactosamine (GalNAc, Tn antigen), as reported by *Helix aspersa* agglutinin (HAA) staining, exclusively in WT Jurkat cells. This may be an unexpected response to the already high-levels of α2-6-sialoglycans on these cells at baseline.

**Figure 4.**
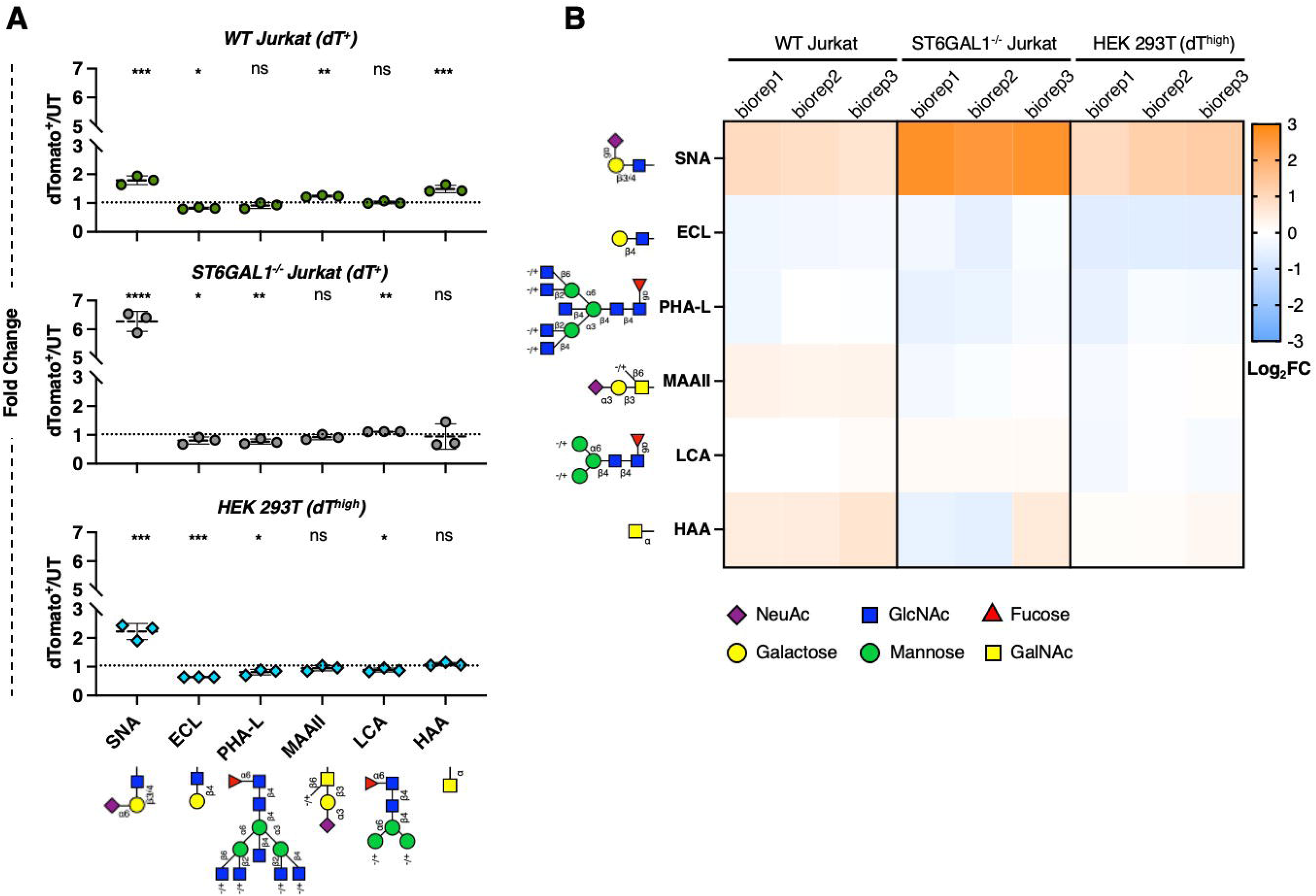
Comparison of changes in glycocalyx composition induced by *ST6GAL1 /dT* mRNA across multiple cell systems. **(A)** Dot plots depicting fold changes in lectin signal across cell systems relative to untransfected (UT) controls 48 hours post-transfection, measured by flow cytometry. Dots represent biological replicates, mean ± SD (*n* = 3). ns = *p* ≥ 0.05, \**p* ≤ 0.05, \*\**p* ≤ 0.01, \*\*\**p* ≤ 0.001, \*\*\*\**p* ≤ 0.0001. Individual controls are shown in Figure 2C, Supplementary Figure S3D-E. One way ANOVA. **(B)** Heat maps depicting Log_2_ Fold Change (Log FC) in lectin signal across model cell systems relative to UT controls 48 hours post-transfection, measured by flow cytometry.

As expected, when comparing the magnitude of the changes, *ST6GAL1*^−/−^ Jurkat cells displayed a more pronounced fold-change in SNA staining upon *ST6GAL1/dT* mRNA delivery (vs. untransfected controls) relative to WT counterparts (**Figure 4B**). Commensurate loss of ECL and PHA-L binding was also more drastic in *ST6GAL1*^−/−^ cells. We also found that HEK 293T cells were more responsive to *ST6GAL1/dT* mRNA compared to WT Jurkat cells, which may be due to their relatively lower baseline α2-6-sialylglycan levels (**Figure 4B, Supplementary Figure S3F**). Similarly, the reduction of ECL and PHA-L staining was more dramatic in HEK 293T cells vs. WT Jurkat cells, suggesting again that different cell types with differing base glycan levels and glycocalyx composition vary in the magnitude of their possible modulation by *ST6GAL1* mRNA.

### Expanding the toolbox and its limitations

To assess whether cellular glycoengineering through mRNA was compatible with sialyltransferases beyond ST6Gal1, we next synthesized mRNAs encoding *ST3GAL4, ST3GAL1,* and *ST6GALNAC4* (**Figure 5A**). ST3Gal4 was chosen as a counterpoint to ST6Gal1 since it also sialylates N-linked glycans but through an α2-3-rather than α2-6- linkage.^43^ In contrast, ST3Gal1 and ST6GalNAc4 act exclusively on O-linked acceptor glycans, producing sialyl- and disialyl-core 1, respectively.^43^

**Figure 5.**
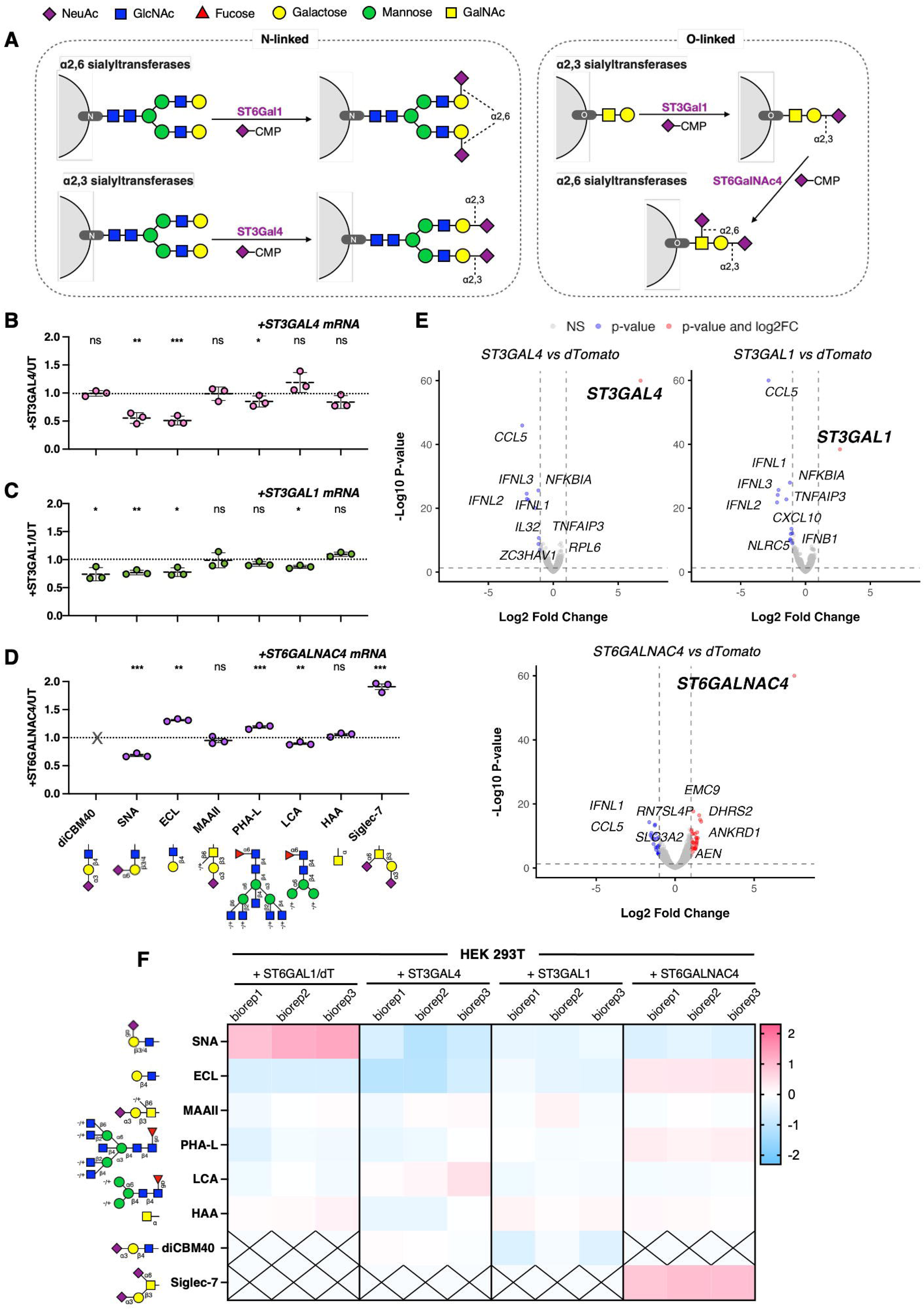
Expanding the mRNA-based sialoglycan engineering toolbox. **(A)** Schematic of sialic acid transfer reactions mediated by ST6Gal1, ST3Gal4, ST3Gal1 and ST6GalNAc4. **(B)** Fold change in lectin binding on HEK 293T cells treated with *ST3GAL4* mRNA relative to untreated control. Individual controls are shown in Supplementary Figure S4B. **(C)** Fold change in lectin binding to HEK 293T cells treated with *ST3GAL1* mRNA relative to untreated control. Individual controls are shown in Supplementary Figure S4D. **(D)** Fold change in lectin binding to HEK 293T cells treated with *ST6GALNAC4* mRNA relative to untreated control. Individual controls are shown in Supplementary Figure S5A. **(E)** RNAseq identifies differentially expressed genes in S*T4GAL4, ST3GAL1,* and *ST6GALNAC4* mRNA versus *dTomato* mRNA transfected HEK 293T cells. The X-axis shows the Log_2_ Fold Change (effect size), and the Y-axis shows the –Log adjusted *p*-values (significance). **(F)** Heatmap displaying lectin binding intensities across *ST6GAL1, ST3GAL4, ST3GAL1,* and *ST6GALNAC4* mRNA treated HEK 293T cells 48 hours post-transfection, measured by flow cytometry. Each row represents an individual lectin, and each column represents a different treatment group, with sub-columns showing biological replicates. Color intensity reflects the Log fold change (Log FC) in lectin binding relative to untreated control, with pink indicating increased binding and blue indicating decreased binding. (**B-D**) Data was collected using flow cytometry. Dots show biological replicates. Mean ± SD (*n* = 3). 48 hours post-transfection. ns = *p* ≥ 0.05, \**p* ≤ 0.05, \*\**p* ≤ 0.01, \*\*\**p* ≤ 0.001. Two-tailed unpaired *t*-test.

We confirmed mRNA delivery to HEK 293T cells using qPCR for all mRNA constructs (**Supplementary Figure S4A**). We expected that increased ST3Gal4 expression would lead to increased staining with the pan-α2-3-sialoglycan selective lectin diCBM40 (dimeric carbohydrate-binding module 40, detecting pan-α2-3-sialyl-linkages);^44^ however, we did not observe any changes after 48 h (**Supplementary Figure S4B**). We verified that diCBM40 was sensitive to α2-3 sialylation in this system by sialidase pre-treatment of HEK 293T cells, which abolished diCBM40 binding as expected (**Supplementary Figure S4C**). Unexpectedly, we detected a substantial decrease in SNA staining (∼50%) 48 h after delivery of *ST3GAL4* mRNA, which suggested that installation of α2-6-sialylglycans was disrupted by ST3Gal4 expression (**Figure 5B**). This may be a result of competition between ST6Gal1 and ST3Gal4 for the same acceptor glycan substrate (asialo LacNAc). In line with this, presentation of asialo LacNAc epitopes as detected by ECL was also decreased; this suggested consumption of open LacNAc acceptors via ST3Gal4-mediated sialylation (**Figure 5B, Supplementary Figure S4B**). Few noteworthy changes were observed for other lectins. Taken together, loss of SNA and ECL staining was indirect evidence that ST3Gal4 was active in cells following mRNA delivery, since it would compete with endogenous LacNAc-processing sialyltransferases (such as ST6Gal1). Our inability to directly detect increased α2-3-sialoglycans may have been the result of limited sensitivity of diCBM40 or production of sialoglycans that could not be recognized by this lectin.

We next evaluated sialyltransferases that act on O-linked glycans. Delivery of mRNA coding for ST3Gal1 to HEK 293T cells was successful as confirmed by qPCR (**Supplementary Figure S4A**). ST3Gal1 overexpression was expected to potentiate the production of sialyl-core 1, which is a preferred binding partner for MAA-II. As with diCBM40 for ST3Gal4, we did not observe a change in MAA-II staining (**Figure 5C**, **Supplementary Figure S4D**). This could be due to MAA-II dominantly recognizing 3-*O*- sulfo-galactose epitopes on HEK 293T cells as an alternative ligand to sialyl-core-1, as previously documented.^27^ To test this, we treated HEK 293T cells with sialidase to remove sialyl T antigen and found that subsequent MAA-II binding was minimally impacted (**Supplementary Figure S4E**). This supported the hypothesis that MAA-II was binding mostly to sulfated galactose in HEK 293T cells instead of sialyl-core-1 since the former would be insensitive to sialidase treatment. For this reason, we concluded that MAA-II was not a useful tool to interrogate ST3Gal1 function in this setting. Beyond our attempts to directly detect the sialoglycan product of ST3Gal1, broad glycan profiling revealed decreases in diCBM40, SNA, and ECL staining following delivery of *ST3GAL1* mRNA (∼30% compared to untransfected control) (**Figure 5C**, **Supplementary Figure S4**). These results were indirect evidence of ST3Gal1 activity on the cell surface, as it may compete with other sialyltransferases that act on core 1 O-linked glycans substrates (such as ST6GalNAc4). Other lectins were minimally impacted.

As with ST6Gal1, we also probed for changes in global sialoglycan content using Ac4ManNAz-labeling following *ST3GAL1* mRNA delivery (**Figure 5C**). These experiments reported no detectable changes in the level of sialic acids installed on HEK 293T cells. Taken together, these data confirmed that not all glycosyltransferases produced the expected on-target glycocalyx edits at levels that were detectable by existing lectins when deployed in mRNA-based glycoengineering workflows. Nevertheless, even these unexpected glycocalyx modifications may be useful in scenarios where transient and broad suppression of sialo- and asialo-LacNAc epitopes is desired.

Finally, we tested mRNA coding for the O-linked α2-6-specific sialyltransferase *ST6GALNAC4* (qPCR in **Supplementary Figure S4A**) for its ability to alter cell surface glycosylation (**Figure 5D**, **Supplementary Figure S5A**). ST6GalNAc4 produces disialyl core 1 which is a preferred binding partner for human Siglec-7.^45^ In line with expectations, transfection with *ST6GALNAC4* mRNA increased staining with Siglec-7- Fc ∼2x over control sample. Surprisingly, we also detected a decrease in SNA staining, which suggested interplay between expression of ST6GalNAc4 and α2-6-sialylation of N-linked glycans, possibly through competition for metabolites. This occurred in concert with increased ECL staining, which also pointed to a reduction in sialylation of LacNAc motifs on N-linked glycans that would normally be processed by ST6Gal1. PHA-L staining was increased, which was consistent with a reduction of N-linked glycan sialylation since sialic acid capping of branched N-linked structures attenuates PHA-L binding. Staining with other lectins including MAA-II, LCA, and HAA was minimally impacted.

Given the lack of expected changes in key lectin binding upon *ST3GAL4* and *ST3GAL1* mRNA delivery, we next performed unbiased transcriptome-wide analysis of sialyltransferase mRNA-transfected cells to test for the potential compensatory expression of ‘off-target’ (endogenous) glycosyltransferases (**Figure 5E, Supplementary Table S3-S5**). Specifically, we were interested to learn whether a reduction in N-linked α2-6-sialylation, as shown by lowered SNA binding, upon ST3Gal4, ST3Gal1, and ST6GalNAc4 overexpression could be the result of downregulation of *ST6GAL1* gene expression as opposed to metabolic competition. We did not observe major changes in any glycosylation-pathway associated genes, including *ST6GAL1* (**Supplementary Figure S5B-D, Supplementary Table S6-S8**). Lowered cellular immune responses compared to control mRNA-transfected cells were observed in all cases, further confirming that our lectin-specific changes were driven by the distinct mRNAs as opposed to mRNA delivery *per se*. This suggested that most changes to other glycans were related to substrate competition as opposed to compensatory up- or downregulation of other sialyltransferases upon transient overexpression.

Taken together, our results confirm that precise glycoengineering outcomes can be realized through delivery of mRNA coding for specific sialyltransferases. While not all encoded sialyltransferases produce the expected effects, select mRNA-based tools provided precise and transient control over mammalian cell glycosylation. This occurred without the need to design and synthesize chemical probes or genetically edit cells.

## Discussion

Delivery of mRNA coding for various sialyltransferases enabled the fast and facile modulation of cell surface sialoglycan composition (**Figure 5F**). *ST6GAL1* mRNA delivery led to an increase of SNA signal, indicating the successful installation of N-linked α2-6-linkages on the surface of mammalian cells. In turn, Siglec-7 confirmed the upregulation of O-linked α-2-6-sialoglycans, at the expense of N-linked residues following *ST6GALNAC4* mRNA delivery. In contrast, we could not directly detect the expected α2-3-sialoglycan products from *ST3GAL4* and *ST3GAL1* mRNA. While increases in α2-3-sialylation were indirectly supported through the reduction of α2-6-linkages and LacNAc acceptor sites, it remains unclear if α2-3-products were indeed produced but at levels below the detection limit of current lectin technologies. Changes to glycan structures unrelated to the specific acceptor or product glycans of these sialyltransferases were mostly minimal. Metabolic labeling with Ac_4_ManNAz confirmed that global sialoglycan content remained similar to previous levels following mRNA delivery (**Supplementary Figure S2D-E**). This suggested that while specific sialoglycan epitopes can be modified, the absolute sialoglycan content of a cell is intrinsically limited.

Precision engineering of the glycocalyx of live cells is a longstanding challenge in glycobiology. Although our results establish this tool’s utility within the context of sialyltransferases, this platform also has broader potential for studying glycosylation and cell surface glycan modifications across a wide range of systems. However, our study also shows that careful validation of glycosyltransferase mRNAs must be performed to fully characterize the breadth of changes that result from potentiating the expression of a given glycosyltransferase.

Our method complements existing strategies for glycoengineering, which permit tracking and modification of cell surface glycans but may carry limitations that constrain their use in certain areas. Genetic engineering such as CRISPR/Cas9-based editing or the genetic integration of additional gene copy numbers induce permanent genomic alterations and generally only allow for on/off expression changes. Moreover, while the SEEL approach can transiently modify glycans directly on a cell surface, it is restricted by substrate availability and labour-intensive glycosyltransferase purification steps. It is also limited in its functionality *in vivo*.^46^ MOE has great potential in labelling and tracking glycan turnover using bioorthogonal chemistry but tends to lack glycan specificity due to metabolic crosstalk.

Since exogenously delivered mRNA is only transiently present in cells, our method allows for fast, dynamic control over glycosylation on a cell surface. For example, we demonstrated that sialylation increased within 24 hours of mRNA delivery and peaked at 24–48 hours in fast dividing cells. Related to this, we also observed a reduction in the glycocalyx edit shortly after mRNA degradation (**Figure 2F**), confirming the reversibility of the mRNA-driven effect. mRNA-mediated overexpression of sialyltransferases would thereby allow for transient glycoengineering.

While careful evaluation of each specific mRNA-encoded sialyltransferase is necessary, the approach can enable programmable control over sialoglycan presentation on human cells *in situ*. It is our expectation that this platform will find utility in dissection of fundamental glycobiology and enable future translational work focused on leveraging glycan-level control over human health.

## Acknowledgements

We are grateful to Dr. Lara Mahal for sharing diCBM40 with us. We thank Sascha Woolcott and Isabelle Da Barp for providing template schemes for glycosylation pathways and Nathalie Simard from the Centre of Immune Analytics (Faculty of Medicine, U of T) for flow cytometry assistance. We thank the Centre for Applied Genomics at SickKids Hospital for Sanger and NGS sequencing and the Cell and Systems Biology Imaging Facility at the University of Toronto. We are also deeply grateful for support and helpful discussions from and with Drs. Obadiah Plante, Michelle Bellerose, and Steven Sazinsky, in the early stages of this project.

## Materials and Methods

### Cell culture

HEK 293T cells were cultured in Dulbecco’s modified Eagle’s medium (DMEM) (Sigma-Aldrich, D5796), supplemented with 10% Fetal bovine serum (FBS)(CORNING, 35-007-CV) and 1% penicillin-streptamycin (Gibco, 15140-122). Cultures were maintained at 37°C in a humidified incubator with 5% CO_2_.

Jurkat cells were maintained in RPMI 1640 medium (Gibco, 11875093) supplemented with 10% heat inactivated FBS, 10 mM HEPES, 0.1 mM NEAA, and 1 mM sodium pyruvate. Cells were maintained at 37°C in a humidified incubator with 5% CO_2_.

Both cell lines were regularly tested for mycoplasma via PCR and were consistently found negative.

### Plasmid design

The coding sequences of *ST6GAL1*, *ST3GAL1*, *ST3GAL4* and *ST6GALNAC4* were PCR-amplified from Jurkat cDNA using Phusion High-Fidelity DNA polymerase (NEB, M0530S). Primers are listed in Table S1. Backbone vector pcDNA3.1(+) (Invitrogen) was linearized with restriction enzymes BamHI-HF (NEB, R3136S) and XhoI (NEB, R0146S). PCR products and linearized plasmid were purified with Monarch Spin PCR & DNA Cleanup Kit (NEB, T1130L), inserts were integrated with Gibson assembly (made in house), and transformed into competent *E. coli* (NEB, C2987H).

ST6GAL1_P2A was synthesized as a gene fragment (Integrated DNA Technologies) for integration into a pcDNA_dTomato vector. The pcDNA_dTomato vector was linearized via PCR, purified, and assembled with inserts via Gibson assembly. A V5 coding sequence was added to pcDNA_ST6GAL1 via PCR, the product was DpnI digested and transformed into *E. coli*. All plasmids were verified with Sanger Sequencing and all primers are listed in **Supplementary Table S9.**

### gRNA design and KO cell generation

A literature validated gRNA sequence targeting ST6GAL1^47^ was cloned into the pSpCas9(BB)-2A-GFP plasmid (Addgene #48138, a kind gift from Feng Zhang). pSpCas9(BB)-2A-GFP was digested using BbsI (NEB, R0539L) and the gRNA sequence was introduced with Gibson assembly. This plasmid was verified with Sanger Sequencing. We transfected this plasmid into wild type Jurkat cells using Lipofectamine 3000 (Invitrogen) according to manufacturer’s instructions and incubated at 37°C with 5% CO_2_ for 48 hours. Single GFP+ cells were FACS sorted into a 96-well tissue culture plate and KO colonies were expanded and validated using Sambucus Nigra Lectin (SNA) staining and flow cytometry. All primers and gRNA sequences are listed in **Supplementary Table S9.**

### *In vitro* transcription

All in vitro transcribed mRNAs were synthesized from a PCR template containing a 5’ T7 promoter and a 3’ 45-nt poly(A) tail. UTRs from pcDNA3.1 were used. mRNA was transcribed using the HiScribe T7 mRNA kit with CleanCap Reagent AG (NEB, E2080S) according to the manufacturer’s instructions. Pseudouridine-modifed mRNA was generated by substituting UTP with Pseudouridine-5’-Triphosphate (NEB, N0433S). Following transcription, reactions were treated with DNase I to remove the template DNA and purified using the Monarch Spin RNA Cleanup Kit (NEB, T2040L). All full mRNA sequences are listed in **Supplementary Table S10.**

### mRNA transfection

HEK 293T and Jurkat cells were plated at 1 x 10^6^ cells per well in 6-well plate, or 2 x 10^5^ cells per 24-well plate. After 24 hours, HEK 293T cells were transfected with mRNA using Lipofectamine MessengerMax (Invitrogen, LMRNA001) following the manufacturer’s manual. Suspension Jurkat cells were immediately transfected once plated. Cells were collected for subsequent analysis 24 hours (Jurkat) or 48 hrs post-transfection (HEK 293T) unless stated otherwise.

### Lectin staining and flow cytometry

HEK 293T and Jurkat cells were collected from their cell culture plates after transfection and washed with PBS. In order not to disturb glycosylated HEK 293T cell surface proteins, cells were not trypsinized, but gently detached from their plates by pipetting. Cells were stained with Zombie NIR or Zombie Violet viability dye (BioLegend) for 10 minutes at room temperature. Then, lectin staining was performed on ice for 20 minutes. Prior to lectin staining, biotinylated lectins were pre-complexed with FITC-streptavidin (BD Biosciences) for 15 minutes on ice in lectin staining buffer (HBSS with Ca^2+^ and Mg^2+^, 1% BSA). For all experiments with FITC and dTomato in the same panel, a BD FACSymphony A3-II instrument with 5 lasers (ultraviolet (349) violet (405 nm), blue (488 nm), yellow-green (561), and red (637 nm)) was used for analysis. All other experiments were conducted on a Cytek Aurora (3 lasers, violet (405 nm), blue (488 nm), and red (640 nm)). Data was analyzed using FlowJo software (V10.10.0, BD Biosciences). Details of lectins and conditions used in lectin staining are listed in **Supplementary Table S12.**

### RNA extraction and RT-qPCR

Total RNA was extracted from cells using a Monarch Total RNA miniprep kit (NEB, T2010S) and reverse transcribed into cDNA using a RevertAid first strand cDNA synthesis kit with oligo(dT)18 primers (Thermo Scientific, K1621). RT-qPCR was performed using PowerUp SYBR green master mix (Applied Biosystems, 3235026) on a Roche LightCycler 480. Results were analyzed using the double delta Ct method. RT-qPCR primers are listed in **Supplementary Table S9.**

### Immunoblotting

After transfection, HEK 293T and Jurkat cells were lysed with 1x RIPA lysis buffer (0.05 M Tris/HCl, pH = 7.4, 0.15 M NaCl, 0.25% deoxycholate, 1% NP-40, 1 mM EDTA) with cOmplete Mini Protease Inhibitor Cocktail (Roche, 11836170001). Lysed cells were spun at 14,000 x g for 10 minutes and the supernatant was collected. Samples were prepared with 5x SDS loading dye and boiled at 95°C for 10 mins.

Proteins were separated by SDS-PAGE using 4-12% Bis-Tris Plus WedgeWell gels (Invitrogen, NW04125BOX) and transferred with iBlot 2 NC Mini Stacks (Invitrogen, IB23002) using a iBlot 2 Dry Blotting system (Invitrogen, IB21001). Membranes were blocked with 5% skim milk in TBS-T for 1 hour. Primary antibodies against V5 (1:2000, Novex, R96025) and GAPDH (1:5000, Cell Signaling Technology, 2118S) were diluted in 3% skim milk in TBST and incubated overnight at 4°C. The next day, membranes were incubated with the corresponding HRP-conjugated secondary antibodies (mouse anti-Rabbit, 1:3000; goat anti-mouse, 1:6000) for 90 minutes at room temperature. Blots were developed using SignalFire™ ECL Reagent (Cell signaling technologies) and imaged with ChemiDoc Imaging Systems (Bio-Rad). Details of antibodies and conditions used in immunoblots are listed in **Supplementary Table S11.**

### Immunofluorescence

HEK 293T cells grown on coverslips (Electron Microscopy Sciences, 12 mm, 1.5) were sterilized in ethanol, rinsed with PBS, and fixed with 4% paraformaldehyde in PBS (Sigma Aldrich, D8537) for 15 minutes at RT. The fixed cells were permeabilized with 0.25% Triton X-100 in PBS for 5 min at room temperature and blocked with 1% BSA (BioShop, 9048-46-8) and 2% goat serum (Corning) in PBS for 1 hour at room temperature. Primary antibodies targeting V5 (Invitrogen, 1:200) were diluted in blocking buffer and incubated with cells at 4°C overnight. The next day, samples were washed 3 times in blocking buffer and incubated with primary antibody against Goglin-97 (Cell Signaling Technologies, 1:150) for 3 hours at RT. Samples were then incubated with fluorescently labelled secondary antibodies (AlexaFlour 488 Goat anti-Rabbit, 1:200, DyLight 650 Goat anti-mouse, 1:200) for 90 minutes at RT in the dark. To visualize F-actin, cells were incubated with DyLight 594 Phalloidin (Cell Signaling Technologies, #12877, 1:300 in PBS) for 20 minutes at RT, followed by DAPI (Roche, 10236276001, 1:500 in PBS) for 15 min at RT. After three PBS washed, coverslips were mounted on glass slides using ProLong Gold Antifade Reagent (Cell Signaling Technologies, #9071S) and left to harden overnight at RT. Samples were sealed with nail polish the next day and stored in 4°C in a dark chamber until imaging. Details of antibodies and conditions used in immunofluorescence are listed in **Supplementary Table S11.**

### Confocal microscopy

Images were acquired on a Leica TCS SP8 confocal microscope (Leica) with 63×/1.40 Oil HC PL APO CS2 objective and LAS X software. Type F Immersion Liquid (Leica) was used. Images were taken at 1024 x 1024 with identical acquisition settings across samples, analyzed with ImageJ/Fiji, and exported as TIFF files.

### Metabolic oligosaccharide engineering

HEK 293T cells were plated at 0.5-1 x 10^5^ cells in 24-well plates. *N*-azidoacetylmannosamine-tetraacylated (Ac_4_ManNAz) was added to each well to a final concentration of 25 µM per well, and HEK 293T cells were transfected with mRNA using Lipofectamine MessengerMax (Invitrogen, LMRNA001) following manufacturer’s instructions. Treatment conditions were varied as specified in **Supplementary Figure S2E.** In all cases, cells were collected and transferred into a 96 U-bottom well plate and washed with DPBS twice to remove residual complete media. Cells were stained with Zombie NIR viability dye and transferred into HBSS buffer containing 1% bovine serum albumin. Next, cells were incubated with biotin-PEG12-dibenzocyclooctyne (1:500 or 1:1000 in HBSS, 1% BSA) for 45 minutes at 37 °C in the dark. Cells were washed twice and subsequently treated with streptavidin-fluorescein tetramers for 25 minutes at 4°C in a dark chamber. Cells were washed twice again and resuspended in a final volume of 200 uL per well in HBSS containing 1% BSA and analyzed on a Cytek Aurora (3 lasers, violet (405 nm), blue (488 nm), and red (640 nm)). Data was analyzed using FlowJo (V10.10.0, BD Biosciences).

### RNA-seq library preparation

Total RNA samples with RNA integrity number scores > 8, as assessed by Bioanalyzer (Agilent), were used for library preparation. Libraries were generated using NEBNext Ultra II RNA Library Prep Kit for Illumina (NEB) in combination with NEBNext Multiplex Oligos for Illumina (Dual Index Primer Set I) (NEB), following the manufacturer’s protocol. Briefly, poly(A) mRNA was isolated from 1 µg of total RNA input using oligo dT magnetic beads, fragmented, and reverse-transcribed to synthesize first-strand cDNA. Second-strand cDNA synthesis was performed to generate double-stranded cDNA. The resulting cDNA product was subjected to end repair/dA-tailing and adaptor ligation. Libraries were PCR-amplified with unique dual index primers for 7 cycles and purified with NEBNext Sample Purification Beads. Libraries were assessed for quality and concentration, and sequencing was performed by the Centre for Applied Genomics (TCAG) at the Hospital for Sick Children (SickKids).

### RNA-seq analysis

>23 million read pairs were sequenced as 150 nt paired end reads for each of three biological replicates by the centre for Applied Genomics (TCAG) using a Novaseq6000 SP 300C. Reads were mapped to Homo_sapiens.GRCh38.114^48^ using STAR aligner 2.7.11b^49^. Counts were obtained using RSubread’s FeatureCount function (Version 2.12.3 or 2.26.0)^50^ and differentially expressed genes were identified with DESeq2 1.38.3 or 1.52.0^51^. Significantly changed gene sets were tested for pathway enrichment with DAVID Knowledgebase v2026_1.

### Statistics

Differentially expressed genes were determined using DESeq2, padj < 0.05. Unpaired Student’s t-test or One-way ANOVA calculated with GraphPad Prism (GraphPad Software Inc.) were used for all other data. Replicates are individually transfected cell passages.

