## Supplementary Figures for "Cell Surface Sialoglycan Engineering Through Exogenous mRNA"

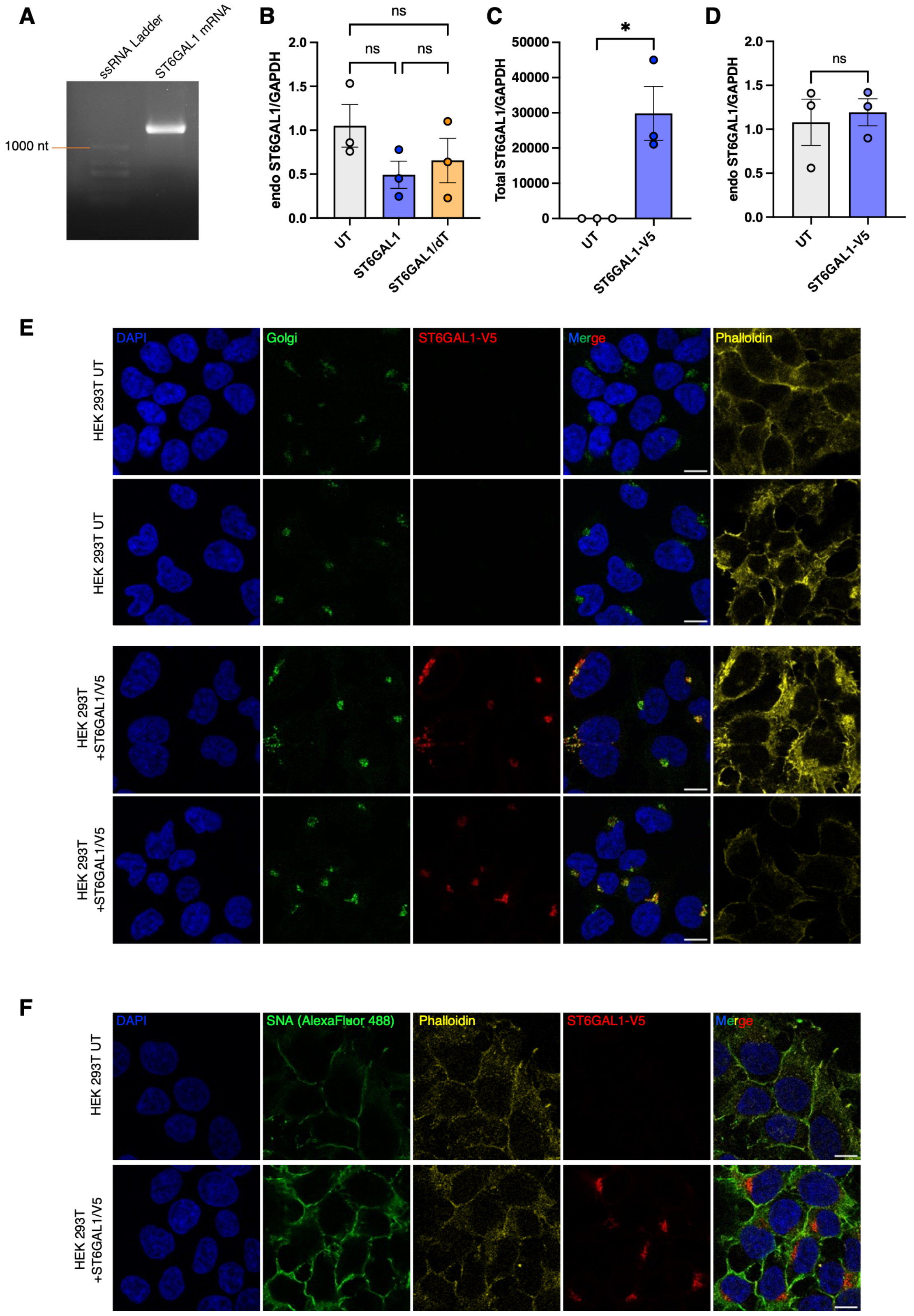

**Supplementary Figure 1. Additional experiments confirming successful *ST6GAL1* mRNA transfection and expression.** (A) Agarose gel to assess *ST6GAL1* mRNA integrity after in vitro transcription and purification. (B) qPCR to quantify endogenous *ST6GAL1* mRNA levels after transfecting HEK 293T cells with *ST6GAL1* and ST6GAL1/dT mRNA. (C) qPCR to quantify total *ST6GAL1* mRNA following *ST6GAL1-V5* mRNA transfection in HEK 293T cells. (D) qPCR to quantify endogenous *ST6GAL1* mRNA expression following *ST6GAL1-V5* mRNA transfection in HEK 293T cells. (E) Additional confocal images of untreated (UT) and *ST6GAL1-V5* transfected HEK 293T cells after immunofluorescence staining for Golgin-97 (green) and ST6Gal1-V5 (red). (F) SNA lectin fluorescence staining to probe the cell surface distribution of α2-6-sialylation on UT and *ST6GAL1-V5* transfected HEK 293T cells.

(B-D) Mean ± SD (n = 3) 48 hours post-transfection. ns = $p \geq 0.05$, *$p \leq 0.05$.
(B) One-way ANOVA followed by Tukey's multiple comparisons test.
(C-D) Two-tailed unpaired t-test.

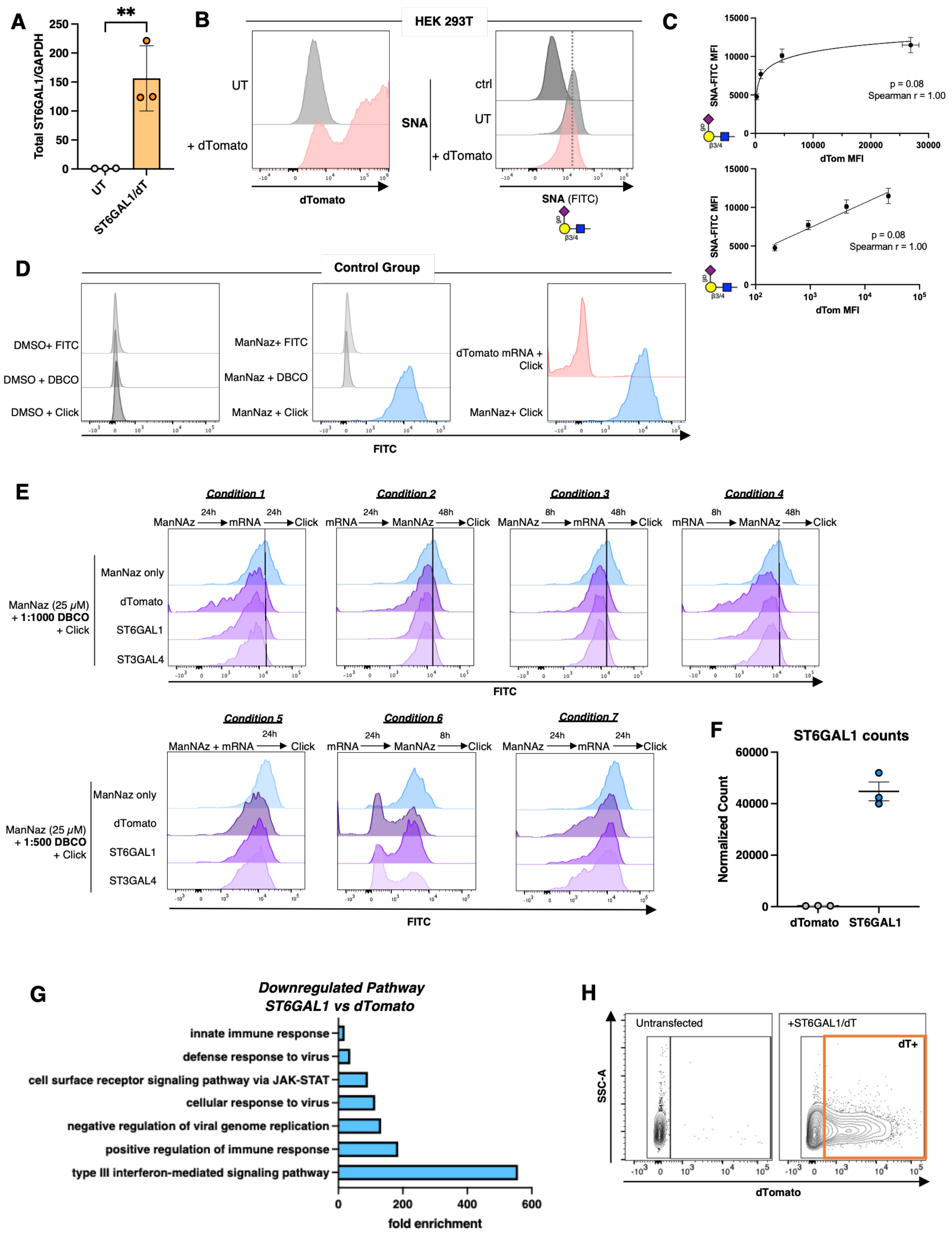
A
Total ST6GAL1/GAPDH
**
UT
ST6GAL1/dT
B
HEK 293T
UT
+ dTomato
dTomato
SNA
ctrl
UT
+ dTomato
SNA (FITC)
C
SNA-FITC MFI
dTom MFI
p = 0.08
Spearman r = 1.00
D
Control Group
DMSO+ FITC
DMSO + DBCO
DMSO + Click
ManNaz+ FITC
ManNaz + DBCO
ManNaz + Click
dTomato mRNA + Click
ManNaz+ Click
FITC
E
Condition 1
ManNAz 24h mRNA 24h Click
Condition 2
mRNA 24h ManNAz 48h Click
Condition 3
ManNAz 8h mRNA 48h Click
Condition 4
mRNA 8h ManNAz 48h Click
ManNaz (25 μM) + 1:1000 DBCO + Click
ManNaz only
dTomato
ST6GAL1
ST3GAL4
FITC
Condition 5
ManNAz + mRNA 24h Click
Condition 6
mRNA 24h ManNAz 8h Click
Condition 7
ManNAz 24h mRNA 24h Click
ManNaz (25 μM) + 1:500 DBCO + Click
F
ST6GAL1 counts
Normalized Count
dTomato
ST6GAL1
G
Downregulated Pathway
ST6GAL1 vs dTomato
innate immune response
defense response to virus
cell surface receptor signaling pathway via JAK-STAT
cellular response to virus
negative regulation of viral genome replication
positive regulation of immune response
type III interferon-mediated signaling pathway
fold enrichment
H
Untransfected
+ST6GAL1/dT
dT+
SSC-A
dTomato

**Supplementary Figure 2. The influence of *ST6GAL1/dT* mRNA delivery on global sialoglycan content and cell transcriptome in HEK 293T cells.** (A) qPCR to assess *ST6GAL1* mRNA expression following *ST6GAL1/dT* mRNA transfection in HEK 293T cells. Mean ± SD (n = 3). **p ≤ 0.01 48 hours post-transfection. Two-tailed unpaired t-test. (B) Left panel − Representative flow cytometry histograms showing dTomato fluorescence from HEK 293T cells transfected with dTomato mRNA after 48 hours. Right panel − SNA staining of HEK 293T cells transfected with dTomato and *ST6GAL1* mRNA after 48 hours. (C) Spearman correlation between dTomato flow cytometry MFI and SNA-FITC flow cytometry MFI intensities in HEK 293T 48 hours post-transfection with *ST6GAL1/dT* mRNA. Upper panel – linear x and y axes. Lower panel – semi-log axes. (D) Control flow cytometry experiments validating the specificity of Ac4ManNAz-mediated sialoglycan click labeling in HEK 293T cells prior to experimental analysis. (E) Global sialoglycan composition in HEK 293T cells across various experimental conditions with Ac4ManNAz and mRNA (see labels), measured with flow cytometry. (F) Normalized read counts obtained from RNAseq of *ST6GAL1* in dTomato control and *ST6GAL1* mRNA transfected cells. Each dot represents a biological replicate. Mean ± SD (n = 3). (G) Significantly downregulated genes (log2 fold change ≥ 1.0) were enriched for genes associated with innate immune responses, virus responses, and interferon-mediated signaling pathways. (H) Gating scheme for Figure 2E.

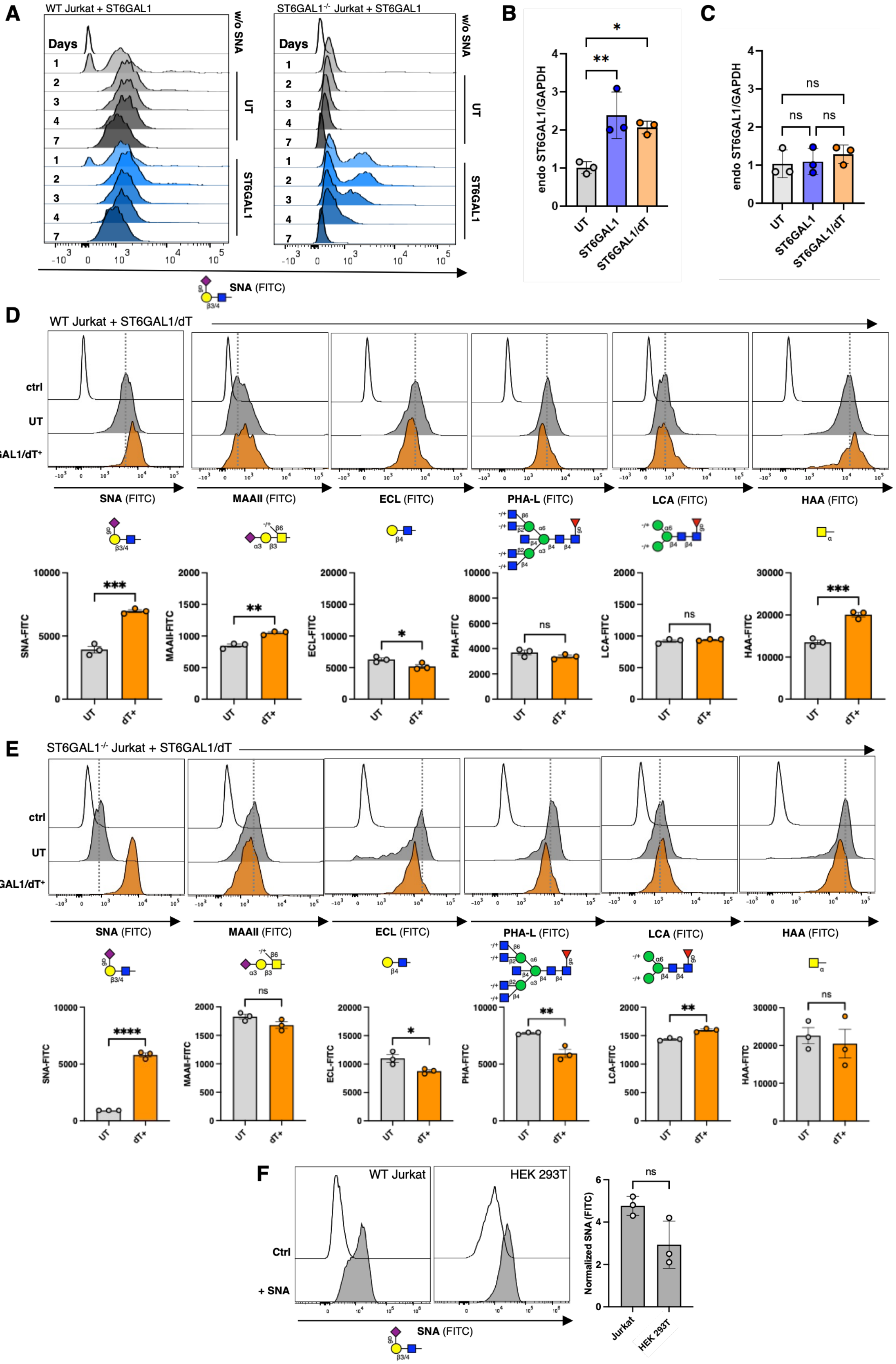

A
WT Jurkat + ST6GAL1
ST6GAL1-/- Jurkat + ST6GAL1
Days
w/o SNA
UT
ST6GAL1
SNA (FITC)
B
endo ST6GAL1/GAPDH
UT
ST6GAL1
ST6GAL1/dT
C
ns
D
WT Jurkat + ST6GAL1/dT
ctrl
UT
ST6GAL1/dT+
SNA (FITC)
MAAII (FITC)
ECL (FITC)
PHA-L (FITC)
LCA (FITC)
HAA (FITC)
SNA-FITC
MAAII-FITC
ECL-FITC
PHA-FITC
LCA-FITC
HAA-FITC
dT+
E
ST6GAL1-/- Jurkat + ST6GAL1/dT
F
WT Jurkat
HEK 293T
Ctrl
+ SNA
Normalized SNA (FITC)
Jurkat
HEK 293T

**Supplementary Figure 3. Impact of *ST6GAL1* and *ST6GAL1/dT* mRNA on global sialoglycan content in Jurkat cells.** (A) Representative flow cytometry histograms of WT and *ST6GAL1*-/- Jurkat cells treated with *ST6GAL1* mRNA after days 1, 2, 3, and 7. (B) qPCR quantifying endogenous *ST6GAL1* transcript levels in untreated (UT), *ST6GAL1* and *ST6GAL1/dT* transfected wild type (WT) Jurkat cells. (C) qPCR quantifying endogenous *ST6GAL1* transcript levels in UT, *ST6GAL1* and *ST6GAL1/dT* transfected *ST6GAL1*-/- Jurkat cells. (D) Representative flow cytometry histograms showing WT Jurkat cells stained with various lectins 24 hours post-transfection with *ST6GAL1/dT* mRNA. Quantification below. (E) Representative flow cytometry histograms showing *ST6GAL1*-/- Jurkat cells stained with various lectins 24 hours post-transfection with *ST6GAL1/dT* mRNA. Quantification below. (F) SNA staining and flow cytometry comparing WT Jurkat and HEK 293T cells. Left panel ‑ SNA staining quantification normalized to unstained control.

(B-F) Mean $\pm$ SD (n = 3), ns = $p \geq 0.05$, *$p \leq 0.05$, **$p \leq 0.01$, ***$p \leq 0.001$, and ****$p \leq 0.0001$

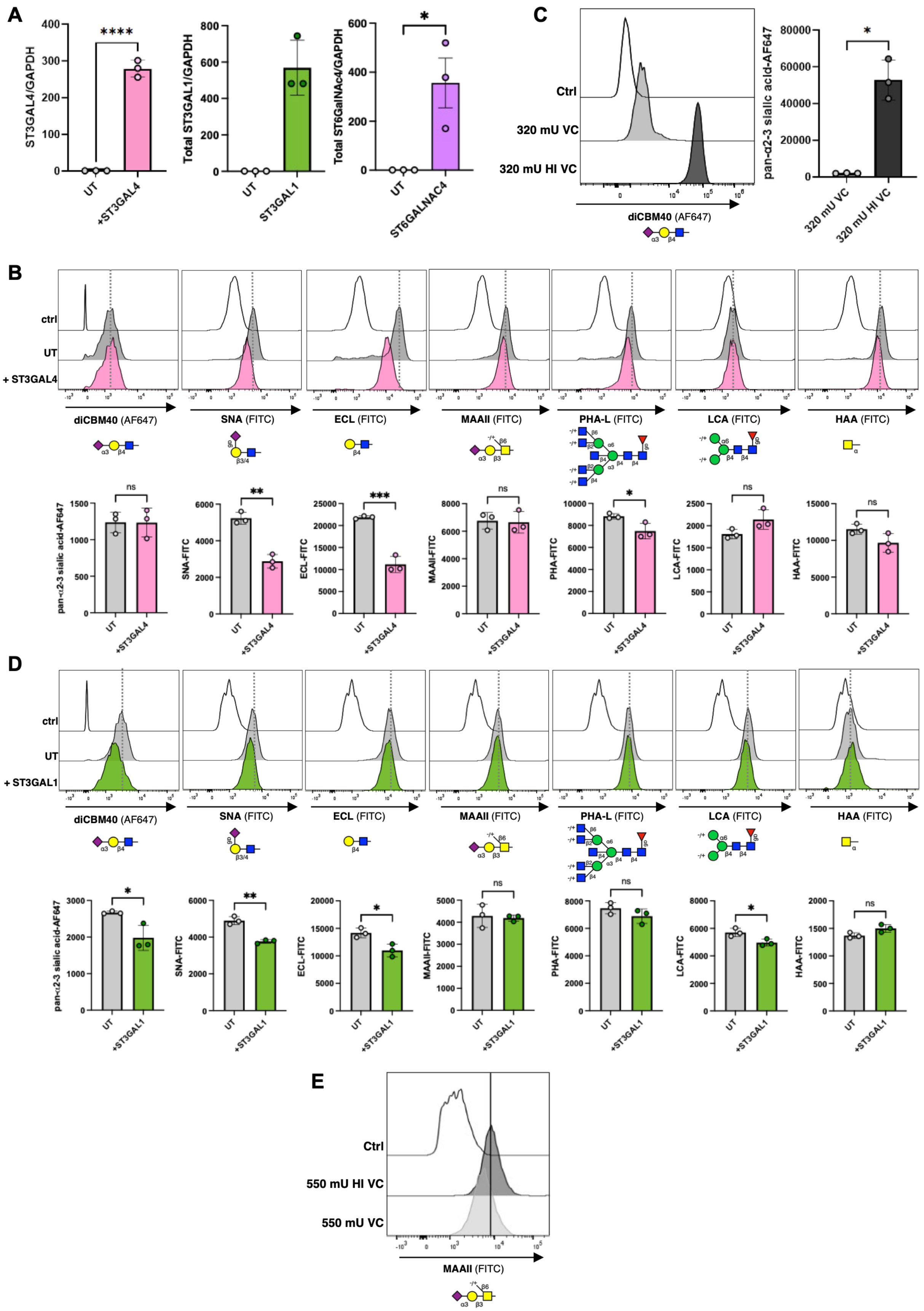

A
ST3GAL4/GAPDH
****
UT
+ST3GAL4
Total ST3GAL1/GAPDH
ST3GAL1
Total ST6GalNAc4/GAPDH
*
ST6GALNAC4
C
Ctrl
320 mU VC
320 mU HI VC
diCBM40 (AF647)
pan-α2-3 sialic acid-AF647
320 mU VC
320 mU HI VC
B
ctrl
UT
+ ST3GAL4
diCBM40 (AF647)
SNA (FITC)
ECL (FITC)
MAAII (FITC)
PHA-L (FITC)
LCA (FITC)
HAA (FITC)
ns
**
***
*
pan-α2-3 sialic acid-AF647
SNA-FITC
ECL-FITC
MAAII-FITC
PHA-FITC
LCA-FITC
HAA-FITC
D
ctrl
UT
+ ST3GAL1
diCBM40 (AF647)
SNA (FITC)
ECL (FITC)
MAAII (FITC)
PHA-L (FITC)
LCA (FITC)
HAA (FITC)
E
Ctrl
550 mU HI VC
550 mU VC
MAAII (FITC)

**Supplementary Figure 4. Impact of other sialyltransferase mRNAs on global sialoglycan content in HEK 293T cells**. (A) qPCR confirming the delivery of *ST3GAL4* (left), *ST3GAL1* (middle), and *ST6GALNAC4* (right) mRNA to HEK 293T cells 48 hours post-transfection. Normalized to GAPDH. (B) Representative flow cytometry histograms of HEK 293T cells stained with various lectins 48 hours post-transfection with *ST3GAL4* mRNA. Quantification below. (C) Validation and quantification of diCBM40 lectin binding to sialoglycans by sialidase treatment. HEK 293T cells post active *V. cholerae* sialidase (VC, light grey) and heat-inactivated (HI VC, dark grey) sialidase treatment, measured in flow cytometry. (D) Representative histograms of HEK 293T cells stained with various lectins 48 hours post-transfection with *ST3GAL1* mRNA. Quantification below. (E) Validation and quantification of MAAII lectin binding to sialoglycans by sialidase treatment. HEK 293T cells post active *V. cholerae* sialidase (VC, dark grey) and heat-inactivated (HI VC, light grey) sialidase treatment, measured in flow cytometry.

(A-D) Mean ± SD (n = 3), ns = $p \geq 0.05$, *$p \leq 0.05$, **$p \leq 0.01$, ***$p \leq 0.001$, and ****$p \leq 0.0001$

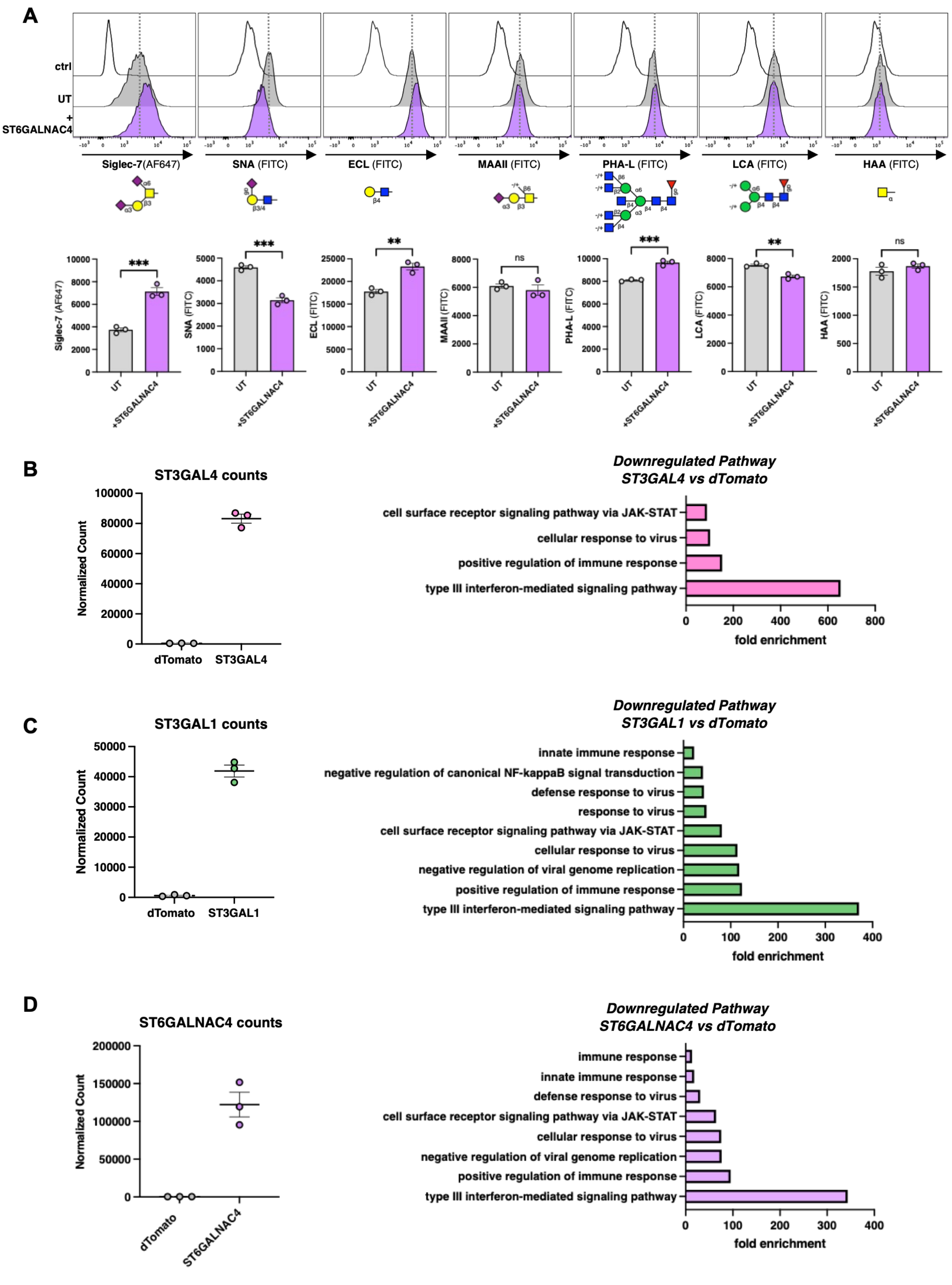
A
ctrl
UT
+ ST6GALNAC4
Siglec-7(AF647)
SNA (FITC)
ECL (FITC)
MAAII (FITC)
PHA-L (FITC)
LCA (FITC)
HAA (FITC)
***
***
**
ns
***
**
ns
Siglec-7 (AF647)
SNA (FITC)
ECL (FITC)
MAAII (FITC)
PHA-L (FITC)
LCA (FITC)
HAA (FITC)
UT
+ST6GALNAC4
B
ST3GAL4 counts
Normalized Count
dTomato
ST3GAL4
Downregulated Pathway
ST3GAL4 vs dTomato
cell surface receptor signaling pathway via JAK-STAT
cellular response to virus
positive regulation of immune response
type III interferon-mediated signaling pathway
fold enrichment
C
ST3GAL1 counts
Normalized Count
dTomato
ST3GAL1
Downregulated Pathway
ST3GAL1 vs dTomato
innate immune response
negative regulation of canonical NF-kappaB signal transduction
defense response to virus
response to virus
cell surface receptor signaling pathway via JAK-STAT
cellular response to virus
negative regulation of viral genome replication
positive regulation of immune response
type III interferon-mediated signaling pathway
fold enrichment
D
ST6GALNAC4 counts
Normalized Count
dTomato
ST6GALNAC4
Downregulated Pathway
ST6GALNAC4 vs dTomato
immune response
innate immune response
defense response to virus
cell surface receptor signaling pathway via JAK-STAT
cellular response to virus
negative regulation of viral genome replication
positive regulation of immune response
type III interferon-mediated signaling pathway
fold enrichment

**Supplementary Figure 5.** (A) Representative flow cytometry histograms for various lectins on HEK 293T cells transfected 48 hours post-transfection with *ST6GALNAC4* mRNA. Quantification below. (B) Left panel − Normalized RNAseq read counts for *ST3GAL4* in dTomato control and *ST3GAL4* transfected cells 48 hours post-transfection. Each dot represents a biological replicate. Mean ± SD (n = 3). Right panel – Significantly downregulated genes (log2 fold change ≥ 1.0) were enriched for genes associated with innate immune responses, virus response, and interferon-mediated signaling pathway. (C) Left panel − Normalized RNAseq read counts for *ST3GAL1* in dTomato control and *ST3GAL1* transfected cells 48 hours-post transfection. Each dot represents a biological replicate. Mean ± SD (n = 3). Right panel – Significantly downregulated genes (log2 fold change ≥ 1.0) were enriched for genes associated with innate immune response, virus response, and interferon-mediated signaling pathways. (D) Left panel − Normalized *ST6GALNAC4* RNAseq read counts in *dTomato* transfected control and *ST6GALNAC4* transfected cells 48 hours-post transfection. Each dot represents a biological replicate. Mean ± SD (n = 3). Right panel – Significantly downregulated genes (log2 fold change ≥ 1.0) were enriched for genes associated with the regulation of immune responses and type III interferon-mediated signaling pathways.

**Supplementary Table legends:**

**Table S1**: Differentially expressed genes in *ST6GAL1* mRNA transfected vs *dTomato* mRNA transfected HEK 293T cells.

**Table S2**: Gene Ontology (GO) analysis of differentially expressed genes in *ST6GAL1* mRNA transfected vs *dTomato* mRNA transfected HEK 293T cells.

**Table S3**: Differentially expressed genes in *ST3GAL4* mRNA transfected vs *dTomato* mRNA transfected HEK 293T cells.

**Table S4**: Differentially expressed genes in *ST3GAL1* mRNA transfected vs *dTomato* mRNA transfected HEK 293T cells.

**Table S5**: Differentially expressed genes in *ST6GALNAC4* mRNA transfected vs *dTomato* mRNA transfected HEK 293T cells.

**Table S6**: Gene Ontology (GO) analysis of differentially expressed genes in *ST3GAL4* mRNA transfected vs *dTomato* mRNA transfected HEK 293T cells.

**Table S7**: Gene Ontology (GO) analysis of differentially expressed genes in *ST3GAL1* mRNA transfected vs *dTomato* mRNA transfected HEK 293T cells.

**Table S8**: Gene Ontology (GO) analysis of differentially expressed genes in *ST6GALNAC4* mRNA transfected vs *dTomato* mRNA transfected HEK 293T cells.

**Table S9**: Oligonucleotides used in this study.

**Table S10**: mRNA sequences used in this study.

**Table S11**: Antibodies used in this study.

**Table S12**: Lectins used in this study.
